# Serum pro-N-cadherin: a biomarker of cardiac fibrosis and diastolic dysfunction in irradiated non-human primates

**DOI:** 10.64898/2026.06.02.728857

**Authors:** Paul D. Ferrell, Drew Neish, Gregory O. Dugan, George W. Schaaf, John D. Olson, Kristianne M. Oristian, Kristofer T. Michalson, Donna Niedzwiecki, Dalane Kitzman, Thomas C. Register, J. Mark Cline, Salvatore V. Pizzo, Chang-Lung Lee

## Abstract

**Background:** The delayed effects of radiation exposure on the heart often manifest as cardiac fibrosis and diastolic dysfunction, which can develop years after exposure. However, no FDA-approved serological biomarker is available to assess the risk of individuals for developing radiation-related heart disease (RRHD).

**Objectives:** Serum pro-N-cadherin (PNC) has shown promise as a marker for predicting the onset of heart failure in the general population. We hypothesize that serum PNC levels will correlate with the risk of RRHD following radiation exposure.

**Methods:** We examined male non-human primates (NHPs) exposed to total-body irradiation (TBI) and unirradiated controls from the Wake Forest University radiation late effects cohort. NHPs exhibited cardiac fibrosis scores ranging from less severe (F0-1) to more severe (F2-3). Cardiac tissue samples collected at necropsy, median 6.8 years post-irradiation, were stained for PNC by immunohistochemistry. PNC was quantified in longitudinal serum samples collected 2, 1 and 0 years before necropsy. The associations of serum PNC levels with cardiac fibrosis scores and echocardiographic parameters were examined.

**Results:** Histological examinations showed aberrant localization of PNC in NHPs with cardiac fibrosis. Elevated serum PNC levels significantly correlated with severe cardiac fibrosis (AUC = 0.81, p = 0.006) and echocardiogram parameters of diastolic dysfunction. Cardiac fibrosis was the only measured comorbidity with a significant difference in serum PNC.

**Conclusions:** Our results demonstrate that serum PNC significantly correlates with cardiac fibrosis and diastolic dysfunction in irradiated NHPs. These findings pave the way for future clinical studies to develop serum PNC as a biomarker of RRHD in humans.

**HIGHLIGHTS:**

- Radiation-related heart disease is an often under-recognized complication of radiation exposure and radiation therapy, which has no FDA-approved biomarkers for assessing risk.
- Our results reveal that serum pro-N-cadherin is a biomarker of cardiac fibrosis and diastolic dysfunction in non-human primates that survived radiation exposure.
- This study lays the foundation for further research into the development of serum pro-N-cadherin as a biomarker for assessing the risk of radiation-related heart disease in humans.

## INTRODUCTION

Radiation-related heart disease (RRHD) is a serious and often under-recognized complication of radiation therapy (RT), particularly in patients treated for cancers located near the chest, such as breast, lung, or mediastinal cancers^1^. For example, while adjuvant RT improves overall survival in patients with invasive breast cancer, survivors who receive RT are at increased risk of late cardiac morbidity and mortality^2–5^. Risk-guided cardioprotection treatments of cancer patients receiving cardiotoxic therapies have shown promise in early-stage clinical trials^6^. Numerous papers reveal a positive association between the risk of heart disease and total radiation doses to the heart and different cardiac substructures^7–10^. Besides cancer patients who receive thoracic RT, epidemiologic studies from atomic bomb survivors also reveal a significant dose-related increase in the risk of cardiovascular disease^11^. For example, one multicenter study reported an estimated excess relative risk per Gy was 14% for heart disease^12^. While much of the focus in radiation-induced cardiovascular injury has historically been on left ventricular (LV) systolic dysfunction, recent research has highlighted the prominence of diastolic dysfunction as a key manifestation of RRHD^13–17^. Diastolic dysfunction refers to the impairment in the heart’s ability to relax and fill properly during the diastolic phase, leading to increased filling pressures and reduced cardiac efficiency, even in the absence of overt systolic dysfunction.

Given that radiation-induced diastolic dysfunction may occur in the absence of noticeable systolic impairment, it presents a unique challenge in diagnosis and management. The pathophysiology of RRHD is multifactorial^18^. Radiation exposure to the heart induces inflammation, vascular injury, and fibrosis, all of which contribute to the stiffening of the left ventricle and impair its ability to relax during diastole^17^. These changes result in altered left ventricular compliance, elevated diastolic pressures, and eventually, the clinical symptoms of heart failure (HF), including shortness of breath, fatigue, and exercise intolerance. Traditional cardiac biomarkers, such as B-type natriuretic peptide (BNP) and troponin, are commonly used to assess HF and monitor cardiac function in a variety of conditions^19^. However, natriuretic peptides and troponin levels measured early after RT have not consistently demonstrated effectiveness in detecting subclinical cardiotoxicity or predicting future cardiomyopathy^1,20,21^. As a result, traditional biomarkers like BNP may not adequately reflect the nuanced and progressive nature of RRHD, especially in cases where diastolic dysfunction is the predominant feature. This limitation underscores the need for more sensitive and specific biomarkers tailored to detect early changes in cardiac function related to RT, as this condition can progress to more severe HF if left unaddressed.

Pro-N-Cadherin (PNC) has emerged as a promising biomarker for predicting the early onset of HF and as an indicator of cardiac fibrosis^22–24^. N-cadherin, a cell adhesion molecule, plays a crucial role in maintaining cardiac tissue integrity and function^25,26^. Under pathological conditions, the defective processing of N-cadherin results in the aberrant localization of the precursor form to cardiac intercalated discs and its release into circulation, which in turn correlates with the extent of fibrosis in the myocardium^22–24^. Thus, elevated levels of PNC reflect ongoing myocardial injury, fibrosis, and remodeling, which are early precursors to HF. Our previous publications indicate that measuring serum PNC levels could help identify patients at risk for developing HF, even before clinical symptoms appear, and may provide a more sensitive marker for cardiac fibrosis compared to traditional biomarkers^22,23^. Common covariates such as age, BMI, and sex were observed to have no significant impact on serum PNC. The goal of this study was to determine the potential of serum PNC for the early detection and monitoring of RRHD in the radiation late effects cohort of non-human primates (NHPs), which had been previously demonstrated to be at risk for RRHD and diastolic dysfunction^27–30^.

## MATERIALS AND METHODS

### The Wake Forest NHP Radiation Late Effects Cohort

The NHPs involved in this study were selected from a larger group of Rhesus macaques (Macaca mulatta) housed at Wake Forest University School of Medicine. All NHPs selected for this study were male. This group is part of The Wake Forest University Nonhuman Primate Radiation Late Effects Cohort (NHP RLEC)^29–32^, which has been built over the past two decades to study the long-term effects of radiation in a large animal model closely related to humans, sharing approximately 93% of their DNA. The RLEC consists of about 200 living animals (both male and female) that are monitored daily and undergo clinical exams, routine blood and fluid collections, and noninvasive imaging such as magnetic resonance imaging (MRI), computed tomography (CT), dual-energy X-ray absorptiometry (DEXA), and ultrasound. The cohort also includes around 130 deceased animals, which were euthanized upon reaching humane endpoints and subsequently underwent full necropsy examinations with the collection of both fixed and frozen tissues, as well as viable cells. All *in vivo* blood collections and post-irradiation procedures were carried out at Wake Forest University School of Medicine, with approval from the university’s Institutional Animal Care and Use Committee (IACUC). The university holds an Assurance on file with the Office for Protection from Research Risks, Office of the Director, National Institutes of Health, which affirms its responsibility for the humane care and use of animals (OPRR #A-3391-01).

Of the animals in the RLEC, approximately 290 were exposed to radiation in various studies at different institutions to investigate radiation effects and medical countermeasures. Around 40 animals were never irradiated and served as controls. The control and irradiated animals were sourced from several institutions, including Wake Forest University, the University of Maryland, the University of Illinois, Armed Forces Radiobiology Research Institute, Lovelace Respiratory Research Institute, Citox Labs, Charles River Laval, and Primate Products. The irradiated animals in this study received TBI doses ranging from 6 to 8.5 Gy under the oversight of the IACUC at their respective institutions. Two irradiation strategies were employed: (1) linear accelerator-derived photons at a nominal mean energy of 2 MeV, delivered at 80 cGy/minute in a split dose, with half delivered anterior-posterior and half posterior-anterior; or (2) Cobalt 60-derived gamma irradiation, delivered bilaterally at 60 cGy/min simultaneously. These doses are potentially lethal: for Rhesus macaques, the LD_10/30_ is approximately 5.5 Gy, the LD_50/30_ is around 6.7 Gy, and the LD_90/30_ is approximately 8 Gy^33^. Surviving animals were subsequently transferred to Wake Forest University School of Medicine for long-term post-radiation monitoring. Previous reports have detailed the irradiation methods, supportive care strategies, and acute effects for many of the animals in this cohort^33–35^. To model North American humans, the RLEC are fed a commercially available diet (5L0P, LabDiet) formulated to model the “typical American Diet” (TAD).

### Ethical Approval

All in vivo blood collections and other post-irradiation procedures were conducted at the Wake Forest University School of Medicine with approval by the Institutional Animal Care and Use Committee of Wake Forest University. Wake Forest University is committed to providing a high-quality program of animal care in compliance with state and federal Animal Welfare Acts and the standards and policies of the US Department of Health and Human Services. Wake Forest University has an Assurance on file in the Office for Protection from Research Risks, Office of the Director, National Institutes of Health, that accepts responsibility for the humane care and use of animals (OPRR #A-3391-01). The Laboratory Animal Care Program of the Wake Forest University School of Medicine complies with the “Principles for Use of Animals”, the “Guide for the Care and Use of Laboratory Animals” [National Research Council. 2011. Guide for the Care and Use of Laboratory Animals: Eighth Edition. Washington, DC: The National Academies Press. https://doi.org/10.17226/12910.)], all provisions of the Animal Welfare Act, and has been accredited by the Association for Assessment and Accreditation of Laboratory Animal Care, International (AAALAC) since April 8, 1966 (AAALAC File #8).

### Echocardiography

NHP were sedated with 10-15 mg/kg ketamine and 0.05-0.15 mg/kg midazolam, and echocardiography was performed using a Logiq S8 ultrasound system (GE Healthcare, Chicago, ILL) by trained NHP ultrasonographers following guidelines for image acquisition and analysis established by the American Society of Echocardiography and European Association of Cardiovascular Imaging^36,37^ combined with recommendations for rhesus macaque echocardiograms reported by Korcarz et al.^38^ as previously described^39,40^. Captured images and video recordings were analyzed using Image-Arena software (TomTec Imaging Systems GMBH, Munich, Germany). Parasternal short and long axis, and apical two (A2C) and four chamber (A4C) views of the heart were obtained to evaluate structure and systolic and diastolic function. Motion (M) mode views at the level of the left ventricle (LV) papillary muscles were captured to assess LV internal diameters at end diastole (LVIDd) and end systole (LVIDs) and fractional shortening (FS). The LV endocardium was traced at end diastole and end systole in A2C and A4C views to obtain end diastolic (EDV) and systolic volumes (ESV) by method of disks. The left atrium (LA) endocardium was traced at end systole in A2C and A4C views to calculate biplane LA end systolic volume (LA Vol BP). Pulsed wave Doppler images taken at the mitral valve inflow tract in the A4C view were used to measure early (E) wave and late (A) wave peak filling velocities, E wave deceleration slope, E wave deceleration time, and the E/A ratio. Tissue Doppler imaging of the lateral annulus of the mitral valve in the A4C view was used to assess early (e’) and late (a’) mitral annular descent velocities, derive the e’/a’ ratio and complete the calculation for the E/e’ ratio, an index of LV filling pressure^41^. The LV outflow tract diameter (LVOT) at the beginning of systole was measured using the parasternal long-axis view. Body surface area (BSA, m^2^) at the time of echocardiogram was calculated as BSA = Body weight^2/3^ × 0.0969, utilizing the animal’s body weight (kg)^42^.

### Pathology and Immunohistochemistry (IHC)

Animals were euthanized and taken to necropsy when they met predefined clinical criteria, such as diagnosis of malignant neoplasia, or in rare cases, died unexpectedly, and a complete necropsy was performed, followed by systematic histologic evaluation of all major organ systems and tissue types. Sections from the right and left ventricular free walls as well as the intraventricular septum were collected from the heart and fixed in 4% neutral buffered formalin for 48 hours and then transferred to 70% ethanol. Hematoxylin and eosin (H&E) slides were routinely prepared and reviewed by board-certified veterinary pathologists. Lung tissue was collected, photographed, and the right and left lungs were weighed individually. Six regions of the lung (left cranial/middle/caudal, right cranial/middle/caudal) were collected from each animal as directed by the supervising veterinary pathologist. Abnormalities such as scarring, gross changes in color, or texture were prioritized for collection. If the lung lobe was normal in appearance, a section was chosen that included a grossly visible bronchus, artery and vein. Six regions of the lung (left cranial/middle/caudal, right cranial/middle/caudal) were fixed in 4% paraformaldehyde and changed to 70% ethanol after 24 h, trimmed and embedded in paraffin. Paraffin-embedded tissue samples were routinely processed and stained with hematoxylin and eosin. Slides from all six sections were reviewed by a board-certified veterinary pathologist and were classified as normal, or mildly, moderately, or severely fibrotic^43^. Cardiac and pulmonary fibrosis was qualitatively scored by the pathologists as none = F0, minimal/mild = F1, moderate = F2, and marked/severe = F3. See **Figure 1** for representative histology of the scoring system. IHC for PNC was performed following the protocol previously described^23^.

**Figure 1.**
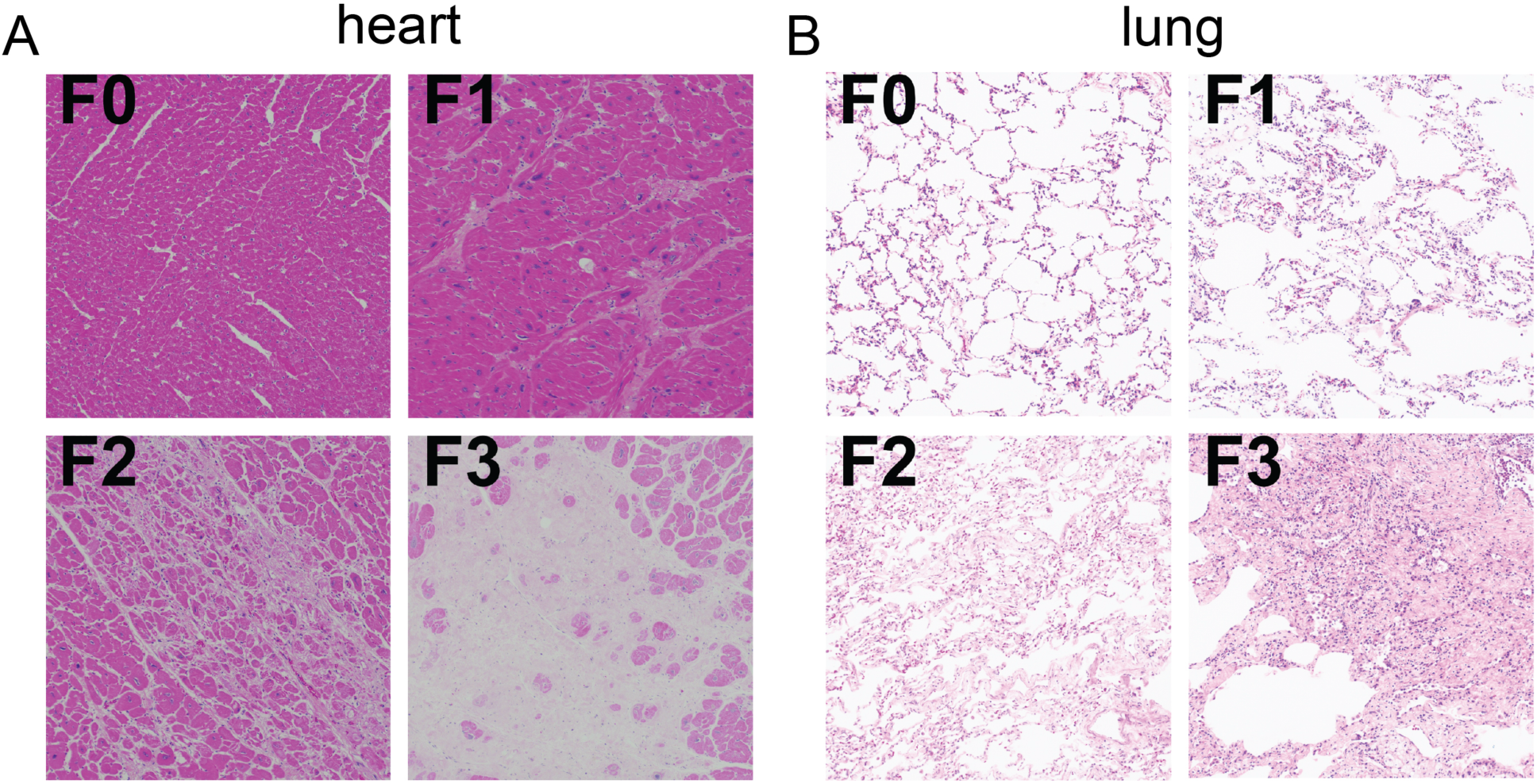
Representative histology of the fibrosis scoring system. (**A**) H&E-stained sections of heart from NHPs at 10x magnification. (F0) Normal myocardial tissue with no evidence of fibrosis. (F1) Minimal/mild myocardial fibrosis – scattered regions of the myocardium contain thin trabeculae of pale-pink staining collagen that separate myocardial bundles. (F2) Moderate myocardial fibrosis – many regions of the myocardium contain variably thick streams of collagen that separate and replace cardiomyocytes. (F3) Marked/severe myocardial fibrosis – significant regions of the myocardium are effaced and replaced by dense accumulations of pale tan mature collagen. (**B**) H&E-stained sections of lung from NHPs at 10x magnification. (F0) Normal pulmonary tissue with no evidence of fibrosis. (F1) Minimal/mild pulmonary fibrosis – scattered regions of the pulmonary interstitium are expanded by thin trabeculae of pale-pink staining collagen. (F2) Moderate pulmonary fibrosis – many regions of the pulmonary interstitium contain variably thick streams of pale-pink collagen. (F3) Marked/severe pulmonary fibrosis – significant regions of the pulmonary architecture are effaced and replaced by dense accumulations of pink, mature collagen.

### Quantification of serum PNC by enzyme-linked immunosorbent assay (ELISA)

The detection and quantification of serum PNC by ELISA were described previously^23^. Recombinant pro domain of N-cadherin, amino acids 26–159 (Accession # AAB22854) was generated and supplied by GenScript (Piscataway, NJ, USA) and used to optimize a PNC sandwich ELISA. High binding ELISA plates (Costar, Kennebunk, ME, USA) were used to bind 1 µg/well of α-PNC antibody 10A10 as the capture antibody. Washes were performed using PBS pH 7.4 0.1% Tween 20. Three washes (300 µL/well) were performed between each of the following steps using Biotek (Winooski, VT, USA) ELx405 Select CW automated plate washer. All steps were performed at room temperature with room temperature equilibrated buffers. Capture antibody was bound overnight at room temperature in PBS pH 7.4 followed by blocking with 300 µL per well blocking buffer 5% non-fat dry milk (Bio-Rad, Hercules, CA, USA) in 1x PBS (Gibco, Grand Island, NE, USA) with 0.1% Tween 20 for 1 h. Serum samples and pro domain analyte standard were applied 100 µL per well in 1% BSA, PBS pH 7.4, 0.1% Tween 20 for 1 h followed by 100 µL/well 1:800 dilution of biotinylated polyclonal sheep α-PNC detection antibody (R&D BAF1388, Minneapolis, MN, USA) in 1% BSA, PBS pH 7.4, 0.1% Tween 20 for 1 h. Streptavidin horseradish peroxidase conjugate (Thermo Fisher, Grand Island, NE, USA) was applied at 100 µL per well, 1:800 in 2% BSA, PBS pH 7.4 for 20 min. Reagent 1-Step Turbo TMB-ELISA (Thermo Scientific, 34022) was substituted for detection following the supplier’s protocol. Plates were read using the Biotek (Winooski, VT, USA) Cytation 3 Imager Reader at an absorbance of 410 nm. Linear dilution of pooled Rhesus Macaque serum was used to verify the cross-reactivity of the assay (**Supplementary Figure 1**).

### Statistical analysis

Descriptive statistics were stratified by irradiation status and expressed as the median and interquartile range (Q1, Q3) for continuous variables, compared across strata with Mann-Whitney U tests, and frequency (N) and percentage (%) for categorical variables, compared across strata with Fisher’s exact tests.

The difference in serum PNC distributions between irradiated and unirradiated NHPs was assessed using non-parametric repeated-measures ANOVA adjusted for timepoint [aligned rank transform (ART) ANOVA] to account for multiple PNC measurements within the same NHP.

The difference in PNC distributions between high cardiac fibrosis (F2-3) and low cardiac fibrosis (F0-1) was evaluated separately within irradiated and non-irradiated NHPs at necropsy, as well as 1 and 2 years prior to necropsy. As cardiac fibrosis was assessed postmortem, fibrosis scores at earlier timepoints were inferred; to avoid biasing a repeated measures framework with this potential measurement error for time points further from necropsy, comparisons between high and low cardiac fibrosis groups were performed cross-sectionally at each timepoint using Mann-Whitney U tests. Samples with any missing longitudinal serum PNC measurements were excluded from this analysis. Receiver operating characteristic (ROC) analysis was performed within irradiated NHPs to determine the association between PNC and high cardiac fibrosis at necropsy. Associations between cardiac fibrosis, irradiation status, and PNC distribution were unadjusted for additional study covariates such as age or BMI, due to concerns for overfitting with small sample sizes, and previous literature suggesting PNC is not significantly associated with age or BMI^22^.

Echocardiogram measures were evaluated across four sub-cohorts: No IR, F0-1; No IR, F2-3; IR, F0-1; IR F2-3. The median (Q1, Q3) is presented for each measure stratified by sub-cohort, and measures were compared across all sub-cohorts using the one-way Kruskal-Wallis test, followed by Dunn’s post-hoc tests. This analysis was conducted separately for all echocardiogram measures 30 months to necropsy, as well as for each NHP’s closest echocardiogram measure within 12 months prior to necropsy. A repeated measures framework was not employed for the 30-month analysis due to sampling intervals and substantial missing data (ex. only 42% of NHPs had the maximum 3 E/e’ measurements).

The relationship between PNC and a range of comorbidities, defined as absence or presence of ever experiencing the comorbidity, was assessed using the Mann-Whitney U test and the median (Q1, Q3) of the PNC distribution was stratified by the absence/presence of each comorbidity. The comorbidities examined were: hypertension; overweight; underweight; tumor (carcinoma, sarcoma, or both); gastrointestinal; heart murmur; lung (defined by abnormal CT density); diabetes; cataracts; hepatic cysts; Kidney (defined by cyst, nephromegaly, abnormal CT density, or abnormal blood urea nitrogen [BUN] levels); arthritis; dermatitis; testicular atrophy; irradiation status; high pulmonary fibrosis; and high cardiac fibrosis. This analysis was limited to PNC at necropsy, to ensure reliability of the cardiac and pulmonary fibrosis scores.

Correlations between PNC and echocardiogram measures were also explored for all measures within 30 months prior to necropsy, as well as the closest measure to necropsy for each NHP within 12 months and 6 months prior to necropsy. Both Pearson and Spearman correlations were calculated to assess both linear and non-linear monotonic relationships. Echocardiogram parameters were considered significantly correlated with serum PNC if either Pearson or Spearman p-value was less than 0.05. Only echocardiogram parameters with significant correlations to PNC were listed in the tables.

All statistical tests were two-sided, with p < 0.05 considered statistically significant. Medians and non-parametric statistical techniques, as detailed above, were selected throughout the analyses for their robustness to small sample sizes and potential deviations from normality.

## RESULTS

### Population dynamics of the NHP cohort

The cohort used in this study included 72 NHPs, 16 unirradiated (No IR) and 56 irradiated (IR), and were selected based on irradiation status, cardiac fibrosis scoring, and sample availability (**Table 1**). Distribution of cardiac fibrosis scores among unirradiated and irradiated NHPs was subject to sample availability and request (**Table 1**). Subjects had a median age (Q1, Q3) of 11.7 (8.8, 16.4) at necropsy. Unirradiated NHPs were significantly older at necropsy than irradiated NHPs (no IR median age (Q1, Q3) = 17.4 (15.4, 19.7), IR median age (Q1, Q3) = 11.3 (8.7, 15.9), p = 0.044). Additionally, unirradiated NHPs had significantly higher calculated BSA relative to irradiated NHPs [no IR median BSA (m^2^) (Q1, Q3) = 0.52 (0.44, 0.60), IR median BSA (m^2^)=0.36 (0.29, 0.44), p < 0.001]. Among IR subjects, the median year since irradiation was 6.8 (5.5-11.0), and the median radiation dose was 6.7 (6.5-7.2) Gy. All irradiated NHPs received TBI.

**Table 1.** Demographic summary of the NHP cohort.

|  | Overall<br>(N = 56) | Unirradiated<br>(N = 10) | Irradiated<br>(N = 46) | p-value |
| --- | --- | --- | --- | --- |
| Age at necropsy<br>(years) |  |  |  |  |
| Median (Q1, Q3) | 11.68 (8.82, 16.44) | 17.40 (15.42, 19.71) | 11.30 (8.67, 15.85) | 0.044 |
| Missing | 6 | 6 | 0 |  |
| BSA (m<sup>2</sup>) |  |  |  |  |
| Median (Q1, Q3) | 0.39 (0.30, 0.48) | 0.52 (0.44, 0.60) | 0.36 (0.29, 0.44) | <0.001 |
| Missing | 0 | 0 | 0 |  |
| IR dose (Gy) |  |  |  |  |
| Median (Q1, Q3) | - | - | 6.74 (6.50, 7.20) | - |
| Missing | - | - | 0 |  |
| Time since<br>irradiation (years) |  |  |  |  |
| Median (Q1, Q3) | - | - | 6.75 (5.48, 11.00) | - |
| Missing | - | - | 0 |  |
| Cardiac fibrosis<br>score, N (%) |  |  |  | 0.053 |
| F0 | 24 (42.9%) | 1 (10.0%) | 23 (50.0%) |  |
| F1 | 20 (35.7%) | 5 (50.0%) | 15 (32.6%) | - |
| F2 | 8 (14.3%) | 3 (30.0%) | 5 (10.9%) | - |
| F3 | 4 (7.1%) | 1 (10.0%) | 3 (6.5%) | - |
| Pulmonary fibrosis<br>score, N (%) |  |  |  | 0.138 |
| F0 | 22 (39.3%) | 7 (70.0%) | 15 (32.6%) |  |
| F1 | 23 (41.1%) | 3 (30.0%) | 20 (43.5%) |  |
| F2 | 9 (16.1%) | 0 (0.0%) | 9 (19.6%) |  |
| F3 | 2 (3.6%) | 0 (0.0%) | 2 (4.3%) |  |

NHP = nonhuman primate, BSA = body surface area, IR = irradiation, F0 = fibrosis score 0, F1 = fibrosis score 1, F2 = fibrosis score 2, F3 = fibrosis score 3. P-values were obtained from Mann-Whitney U tests to compare continuous variables, and from Fisher's exact test for the categorical cardiac fibrosis score, across irradiation groups. Results are restricted to only NHPs with complete longitudinal serum PNC data.

Cardiac and pulmonary fibrosis were scored by board-certified pathologists using a 0 to 3 scale (F0 to F3), with F3 being the most severe (**Figure 1, A and B**). To examine the cardiac function of NHPs that exhibited low (F0-1) versus high (F2-3) cardiac fibrosis scores, we analyzed echocardiogram parameters among four sub-cohorts (No IR F0-1, No IR F2-3, IR F0-1, and IR F2-3) (**Figure 2, A to H, Supplementary Tables 1 and 2**). The ratio of mitral valve peak velocity to left ventricle peak tissue velocity E-Wave (E/e’ lateral) was assessed within 30 months before necropsy and was significantly higher in IR cardiac F2-3 NHPs compared with any of the sub-cohorts (IR F2-3 vs. IR F0-1, p<0.05; IR F2-3 vs. No IR F2-3, p<0.01; IR F2-3 vs. No IR F0-1, p<0.001) (**Figure 2C**). The value of E/e’ lateral assessed within 12 months before necropsy was also significantly higher in IR cardiac F2-3 NHPs compared to IR cardiac F0-1 NHPs (**Figure 2F**). In addition, the left atrium diameter at systole (LA diameter systole) was significantly increased in IR cardiac F2-3 NHPs compared with IR cardiac F0-1 NHPs (p<0.05) within 30 and 12 months before necropsy, respectively (**Figure 2, D and G**). Moreover, IR cardiac F2-3 NHPs exhibited a significant increase in the left atrium diameter to aortic root diameter–systole (LA/AO systole) compared to IR cardiac F0-1 NHPs within 30 and 12 months before necropsy, respectively (**Figure 2, E and H**). Ejection fractions within 30 months prior to necropsy were significantly higher in the IR cardiac F2-3 NHPs relative to the No IR cardiac F2-3 group (**Figure 2A and B**). Together, these results indicate that high cardiac fibrosis scores of IR NHPs correlate with an increase in diastolic left ventricular filling pressure and diastolic dysfunction.

**Figure 2.**
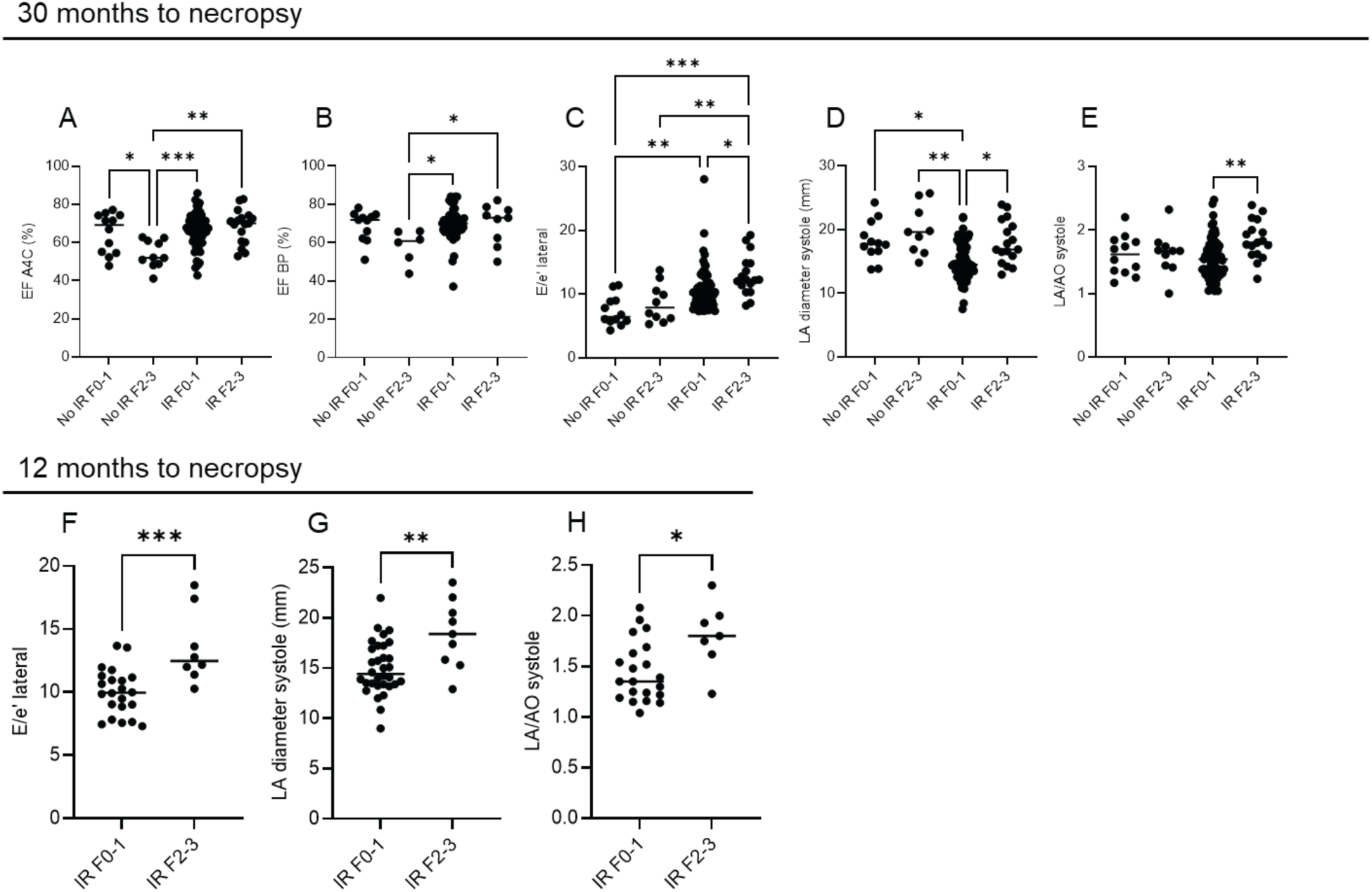
Echocardiogram parameters by sub-cohort. (**A**) EF A4C = Left ventricular ejection fraction apical 4-chamber, (**B**) EF BP = Left ventricular ejection fraction biplane, (**C**) E/e’ lateral = ratio of mitral valve peak velocity to left ventricle peak tissue velocity E-Wave, (**D**) LA diameter systole = Left atrium diameter systole – antero-posterior, (**E**) LA/AO systole = Left atrium diameter to aortic root diameter – systole. A-E dots represent available echocardiogram examinations with corresponding serum samples drawn within 6 months of examination over 0-2 years prior to necropsy (30 months to necropsy) and are analyzed by nonparametric one-way Kruskal-Wallis test with Dunn’s post-hoc tests. (**F**) E/e’ lateral = ratio of mitral valve peak velocity to left ventricle peak tissue velocity E-Wave, (**G**) LA diameter systole = Left atrium diameter systole – antero-posterior, (**H**) LA/AO systole = Left atrium diameter to aortic root diameter – systole. F-H dots represent individual NHPs and are analyzed by Mann-Whitney U test. Horizontal line represents median value. Insufficient unirradiated NHP echocardiogram examinations within 12 months to necropsy for analysis. (* p < 0.05; ** p < 0.01; *** p < 0.001).

### Elevated serum PNC in irradiated NHP correlates with increased cardiac fibrosis

To examine the expression of PNC in the heart of NHPs, cardiac tissues collected post-necropsy and independently scored for fibrosis were stained for PNC by IHC. Our results showed that the hearts of unirradiated NHPs without fibrosis (F0) exhibited perinuclear staining of PNC (**Figure 3A**). In contrast, PNC^+^ cells were substantially enriched in the hearts of unirradiated and irradiated NHPs that exhibited severe cardiac fibrosis (F3) (**Figure 3, B and C**). PNC IHC staining in fibrotic cardiac myocytes was present in a patchy cytoplasmic distribution, most intense at intercalated discs (**Figure 3B, top panel**) and hypercontraction bands (**Figure 3C, top panel**). Cardiac myocytes of affected animals showed other pathologies, including variation in myofiber diameter, and karyomegaly, evident in panels **Figure 3B** and **Figure 3C**. The patterns of aberrant localization of PNC are consistent with PNC localization observed in failing human hearts^23^.

**Figure 3.**
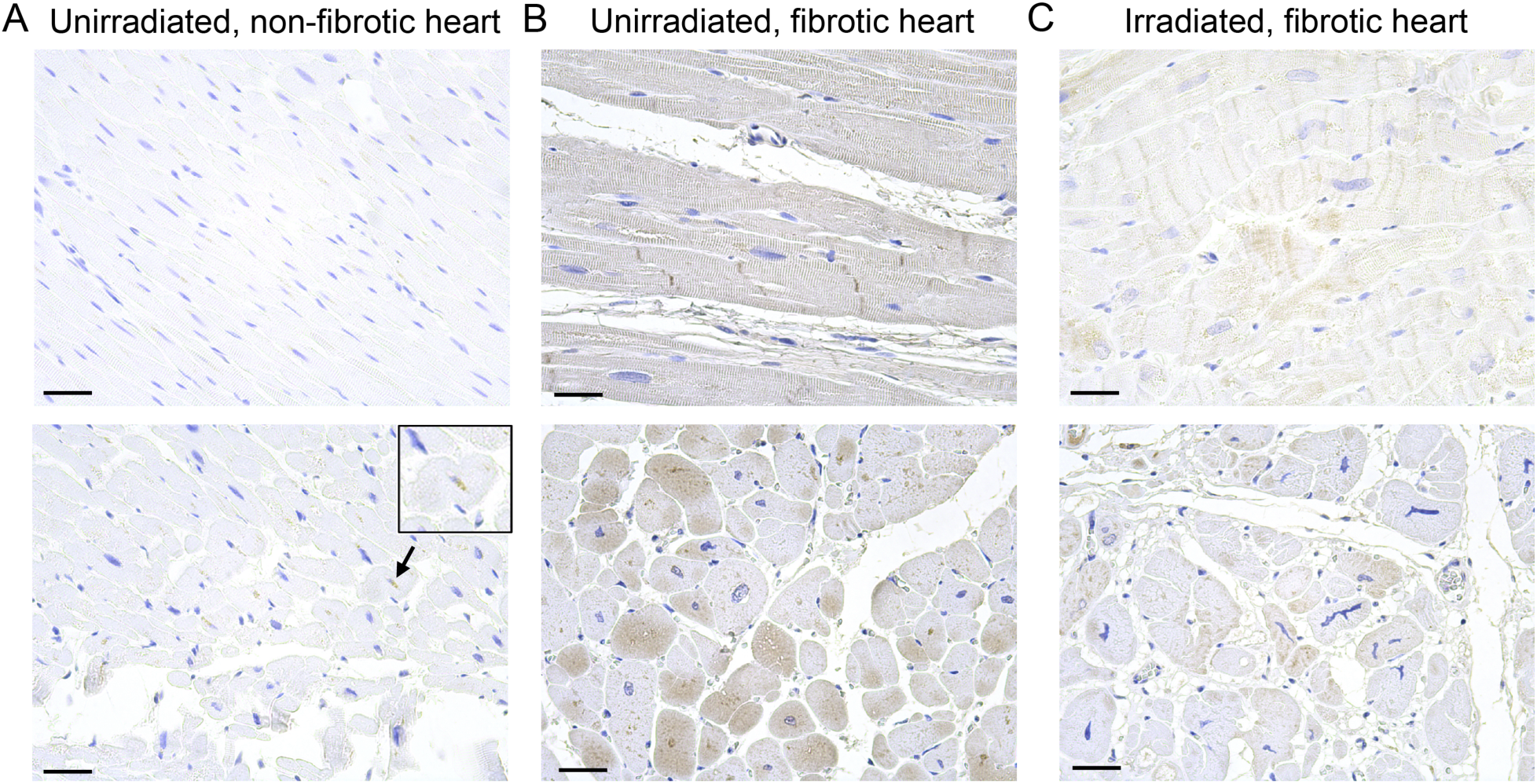
Aberrant localization of PNC in irradiated and unirradiated NHP fibrotic cardiac tissue. Representative images of PNC staining (brown stain) of the hearts at necropsy from (**A**) unirradiated NHPs without cardiac fibrosis (F0), (**B**) unirradiated NHPs with cardiac fibrosis (F3) and (**C**) irradiated NHPs with cardiac fibrosis (F3). The top images are longitudinal sections, and the bottom images are transverse sections. Each image represents an individual NHP. In non-fibrotic cardiac tissues, perinuclear staining of PNC, consistent with normal N-cadherin processing, was observed (black arrow and inset zoomed image). In contrast, PNC IHC staining in fibrotic cardiac myocytes was present in a patchy cytoplasmic distribution, most intense at intercalated discs (B, top panel) and hypercontraction bands (C, top panel). Cardiac myocytes of affected animals showed other pathologies, including variation in myofiber diameter and karyomegaly, evident in panels B and C. Scale bars are 25µm.

Longitudinal serum samples 2, 1, and 0 years prior to necropsy were available from 11 unirradiated and 46 irradiated NHPs. Median serum PNC was higher in no IR cardiac F2-3 NHPs compared with no IR cardiac F0-1 NHPs at necropsy, although the difference was not statistically significant (**Figure 4A**). Serum PNC increased significantly (p=0.005) in irradiated NHPs with more severe cardiac fibrosis, F2-3, at necropsy relative to irradiated NHPs with less severe cardiac fibrosis, F0-1 (**Figure 4B**). ROC analysis at necropsy of serum PNC demonstrated a significant discrepancy between irradiated NHPs with cardiac fibrosis scores F2-3 relative to irradiated NHPs with cardiac fibrosis scores F0-1 (AUC = 0.81, p = 0.006) (**Figure 4C**). No significant difference was observed between irradiated NHPs with pulmonary fibrosis scores F2-3 versus pulmonary fibrosis scores F0-1 at necropsy (**Figure 4, D and E**). No severe pulmonary fibrosis (F2-3) was observed in unirradiated NHPs. Collectively, our findings support the hypothesis that serum PNC is a marker of cardiac fibrosis in irradiated NHPs.

**Figure 4.**
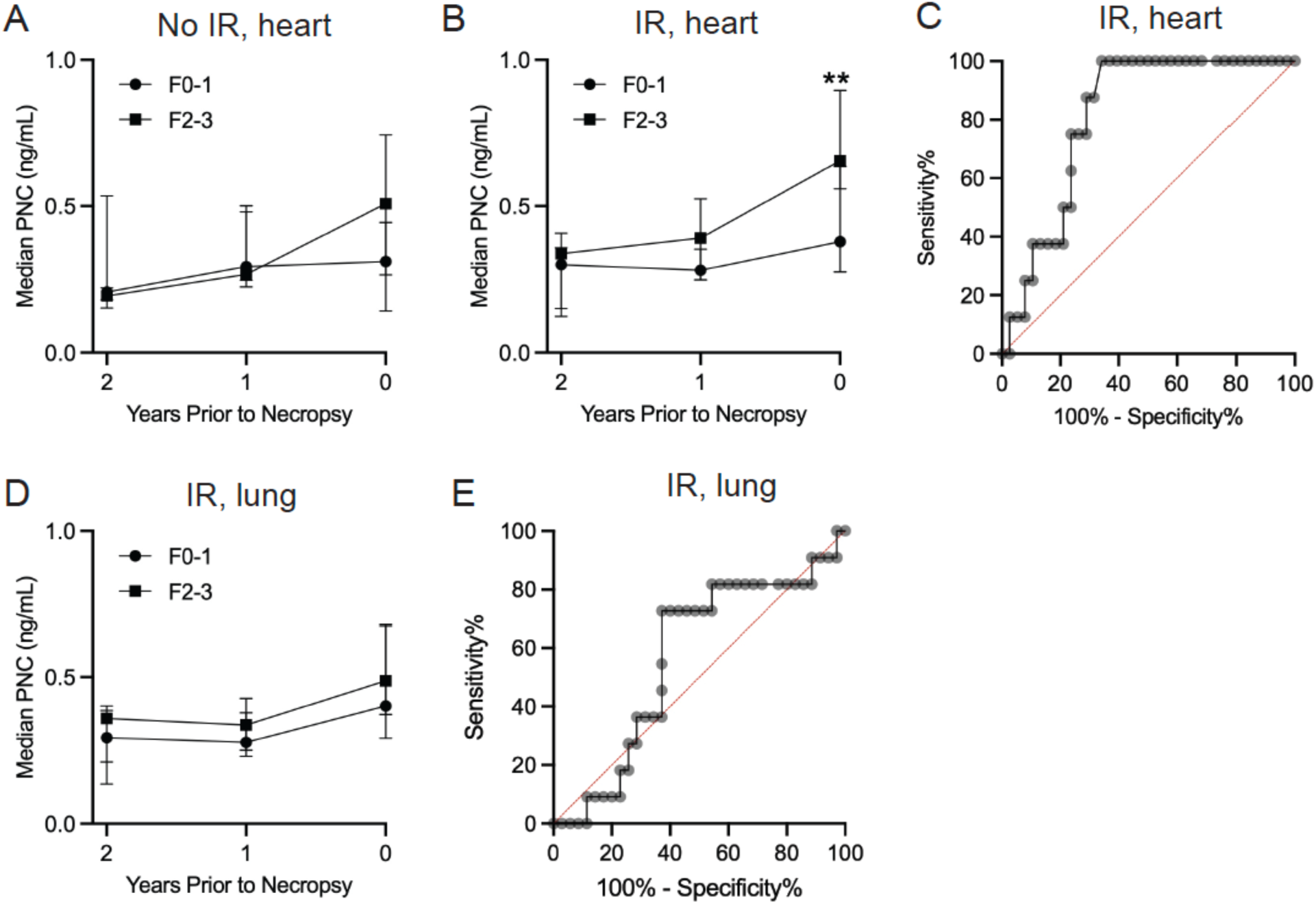
Elevated serum PNC in irradiated NHPs with cardiac fibrosis. (**A**) Longitudinal serum median PNC levels of unirradiated NHPs at 2-years, 1 year, and 0-year prior to necropsy stratified by cardiac fibrosis score at necropsy (F0-1 (N) = 6, F2-3 (N) = 4), error bars represent interquartile range. These differences were not significantly different by Mann-Whitney U test. (**B**) Longitudinal serum median PNC levels of irradiated NHPs at 2-years, 1 year, and 0-year prior to necropsy stratified by cardiac fibrosis score at necropsy (F0-1 (N) = 38, F2-3 (N) = 8), error bars represent interquartile range. Irradiated NHP with cardiac fibrosis scores F2-3 have significantly higher serum PNC compared to irradiated NHPs with cardiac fibrosis scores F0-1 at necropsy by Mann-Whitney U test (p=0.005). (**C**) ROC analysis of irradiated NHPs with cardiac fibrosis scores F0-1 relative to F2-3 at necropsy (IR F0-1, n=38; IR F2-3, n=8; AUC=0.81; [95% CI, 0.69-0.93]; p=0.006). (**D**) Longitudinal serum median PNC levels of irradiated NHPs at 2-years, 1 year, and 0-year prior to necropsy stratified by pulmonary fibrosis score at necropsy (F0-1 (N) = 35, F2-3 (N) = 11), error bars represent interquartile range. These differences were not significantly different by Mann-Whitney U test. (E) ROC analysis of irradiated NHPs with pulmonary fibrosis scores F0-1 relative to F2-3 at necropsy (IR F0-1, n=35; IR F2-3, n=11; AUC=0.57; [95% CI, 0.38-0.76]; p=0.511). (* p < 0.05; ** p < 0.01).

### Serum PNC correlates with echocardiogram parameters of diastolic function

To determine the correlation between serum PNC and echocardiogram parameters, we examined echocardiogram measurements performed within 6 months of serum collection. The distribution of time of echocardiograms before necropsy was not significantly different between any of the sub-cohorts (**Supplementary Figure 2**). Serum PNC levels were significantly correlated with multiple NHP echocardiogram parameters of diastolic function (**Table 2**), including left ventricle major axis length diastolic difference, lateral mitral annulus a’, septal mitral annulus a’, lateral mitral annulus e’, lateral mitral annulus e’/a’, lateral mitral annulus E/e’, mitral valve E-wave peak velocity, mitral valve antegraded flow E/A, left atrial volume A2C – end systole, left atrial volume BP – end systole, and left atrial area A2C – end systole. Since the E/A ratio is a non-monotonic scale for the measure of diastolic dysfunction, additional correlations with lower and upper E/A quartiles are performed (Q1-2, Q3-4). As time nears necropsy, PNC correlations with echocardiogram parameters became greater, and additional echocardiogram parameters became significantly correlated: Aortic valve peak gradient, aortic valve mean gradient, aortic valve peak velocity, aortic valve mean velocity, aortic valve velocity time integral, and left ventricular ejection fraction A2C (**Tables 3 and4).**

**Table 2.**
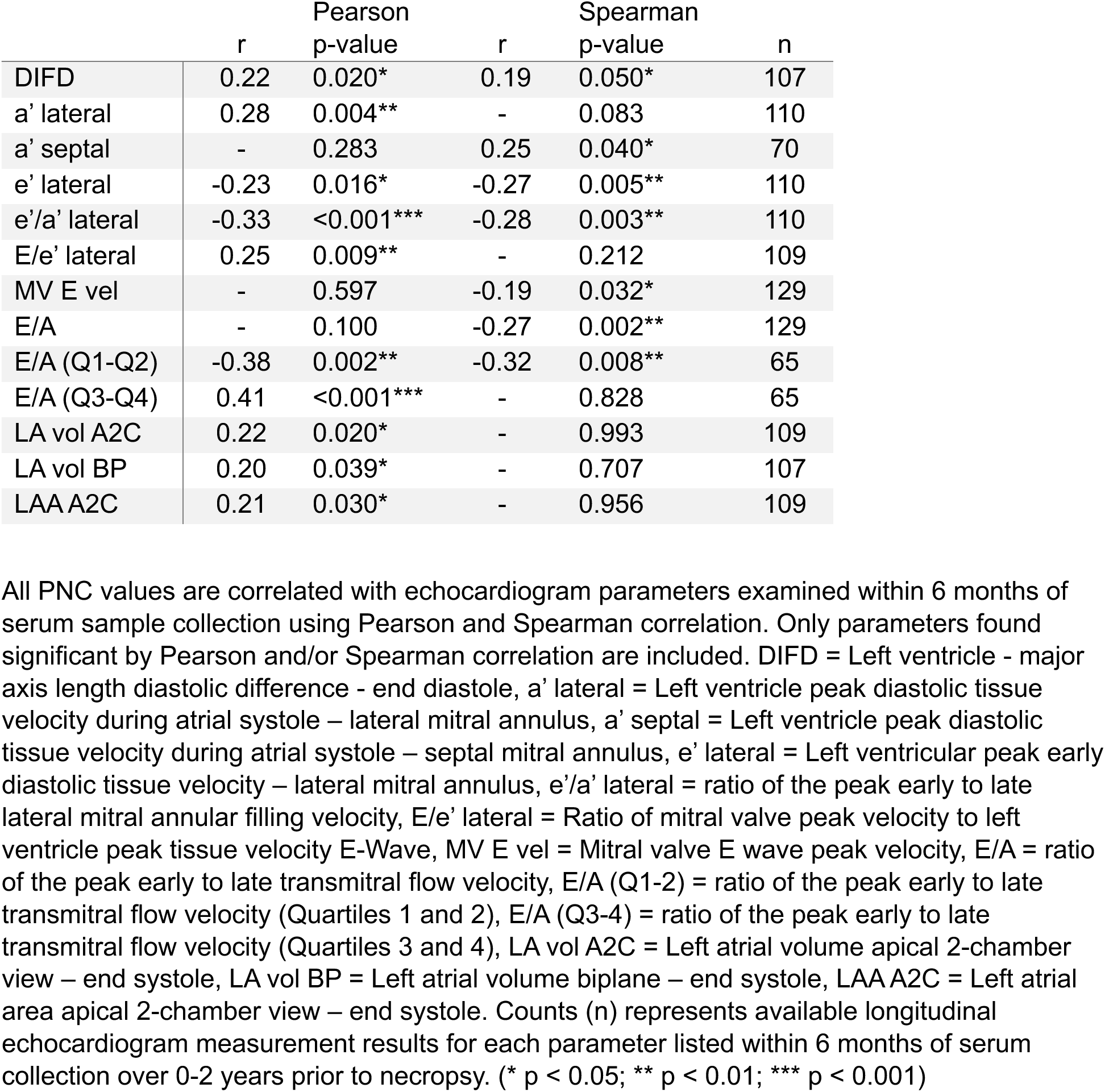
Summary of echocardiogram parameter correlations with PNC 30 months to necropsy.

**Table 3.**
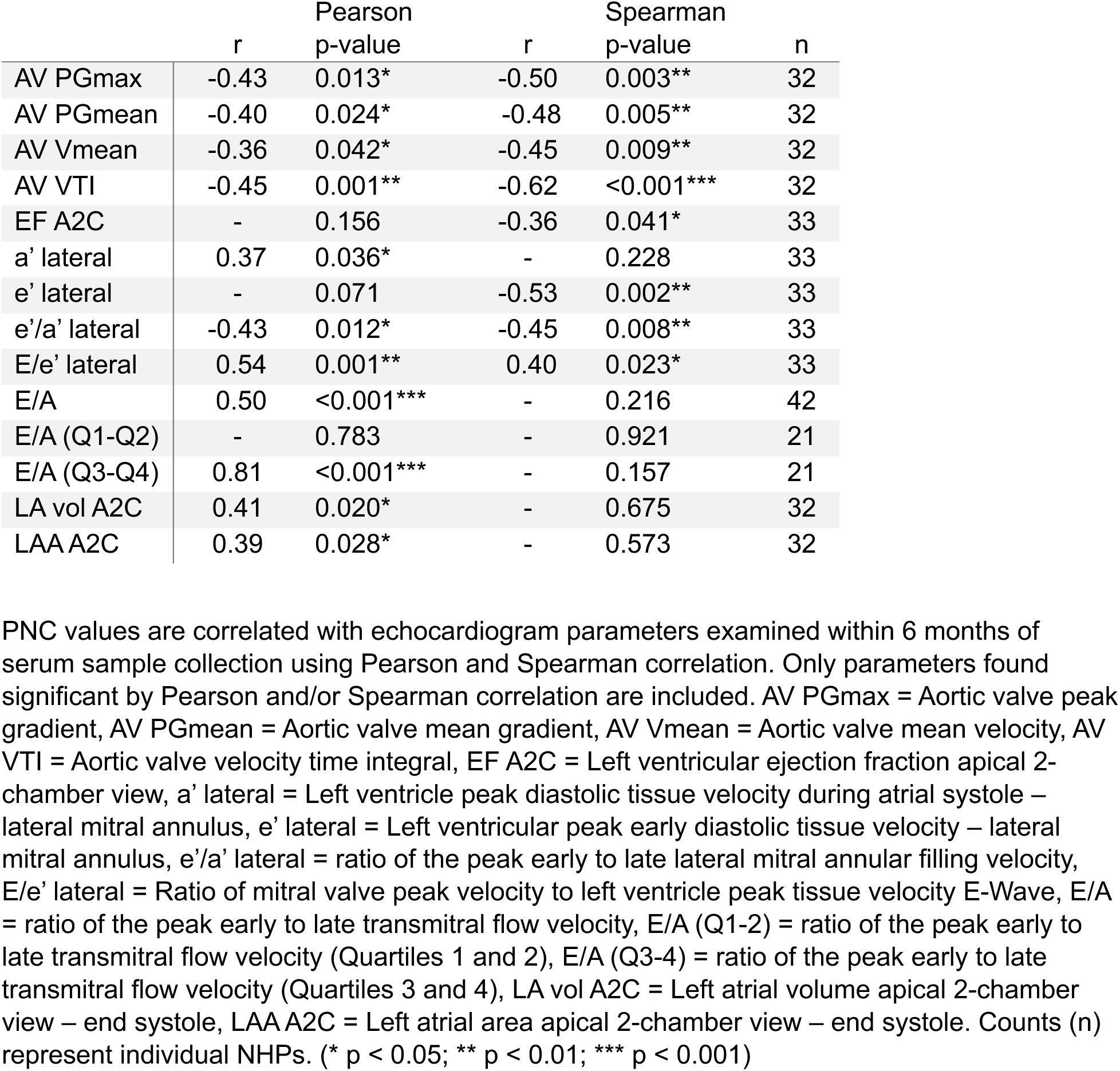
Summary of echocardiogram parameter correlations with PNC 12 months to necropsy.

**Table 4.** Summary of echocardiogram parameter correlations with PNC 6 months to necropsy.

|  | r | Pearson<br>p-value | r | Spearman<br>p-value | n |
| --- | --- | --- | --- | --- | --- |
| AV PGmax | -0.50 | 0.014* | -0.62 | 0.001** | 24 |
| **AV PGMEAN** | -0.43 | 0.036* | -0.55 | 0.006** | 24 |
| **AV VMEAN** | - | 0.065 | -0.50 | 0.013* | 24 |
| AV Vmax | - | 0.075 | -0.45 | 0.013* | 30 |
| AV VTI | -0.48 | 0.016* | -0.72 | <0.001*** | 24 |
| **EF A2C** | - | 0.124 | -0.47 | 0.017* | 25 |
| **E' LATERAL** | - | 0.114 | -0.57 | 0.003** | 25 |
| **E'/A' LATERAL** | -0.44 | 0.029* | -0.46 | 0.020* | 25 |
| **E/E' LATERAL** | 0.56 | 0.004** | 0.43 | 0.032* | 25 |
| **E/A** | 0.49 | 0.006** | - | 0.072 | 30 |
| E/A (Q1-Q2) | - | 0.230 | - | 0.314 | 15 |
| E/A (Q3-Q4) | 0.84 | <0.001*** | - | 0.217 | 15 |
| LA vol A2C | 0.52 | 0.009** | - | 0.784 | 24 |
| LA vol BP | 0.43 | 0.041* | - | 0.801 | 22 |
| LAA A2C | 0.48 | 0.018* | - | 0.673 | 24 |

PNC values are correlated with echocardiogram parameters examined within 6 months of serum sample collection using Pearson and Spearman correlation. Only parameters found significant by Pearson and/or Spearman correlation are included. AV PGmax = Aortic valve peak gradient, AV PGmean = Aortic valve mean gradient, AV Vmean = Aortic valve mean velocity, AV VTI = Aortic valve velocity time integral, EF A2C = Left ventricular ejection fraction apical 2-chamber view, e' lateral = Left ventricular peak early diastolic tissue velocity – lateral mitral annulus, e'/a' lateral = ratio of the peak early to late lateral mitral annular filling velocity, E/e' lateral = Ratio of mitral valve peak velocity to left ventricle peak tissue velocity E-Wave, E/A = ratio of the peak early to late transmitral flow velocity, E/A (Q1-2) = ratio of the peak early to late transmitral flow velocity (Quartiles 1 and 2), E/A (Q3-4) = ratio of the peak early to late transmitral flow velocity (Quartiles 3 and 4), LA vol A2C = Left atrial volume apical 2-chamber view – end systole, LA vol BP = Left atrial volume biplane – end systole, LAA A2C = Left atrial area apical 2-chamber view – end systole. Counts (n) represent individual NHPs. (\* p < 0.05; \*\* p < 0.01; \*\*\* p < 0.001)

To correct for the potential impact of BSA and age on echocardiogram measures^44^, all measures with significant differences by sub-cohort were additionally explored with multivariable regressions adjusting for these covariates (**Supplementary Table 3**). In our analyses (**Tables 2 and4**), we observed that key echocardiographic measures were significantly associated with both high cardiac fibrosis and serum PNC. However, due to concerns about statistical power resulting from a limited sample size, we employed unadjusted non-parametric tests to examine these associations, which did not account for age or body surface area (BSA), potential confounders commonly considered in echocardiographic analyses.

To account for the effects of age and BSA on echocardiographic measures, in **Supplementary Table 3**, we examined the associations between these measures and high cardiac fibrosis scores and serum PNC, within a linear regression framework that included adjustment for age and BSA. As shown in this table, E/e′ lateral, LA diameter systole, and LA/AO ratio in systole remained significantly associated with high cardiac fibrosis even after adjustment for age and BSA. We also found that E/e′ lateral was significantly positively associated with serum PNC after adjustment. While AV PGMax showed a significant inverse association with serum PNC in the unadjusted model, the statistical significance was not retained after adjustment.

Together, these findings suggest that while serum PNC was not associated with all echocardiographic measures, its association with E/e′ lateral was comparable to the association between E/e’ lateral and high cardiac fibrosis score, with both relationships demonstrating similar effect sizes and statistical significance. Importantly, the association between serum PNC and E/e′ lateral remained after adjustment for age and BSA, highlighting the potential value of PNC as a serum biomarker of diastolic function.

### Correlations of serum PNC with comorbidities of irradiated NHPs

Criteria for the diagnosis of comorbidities have previously been reported^45^. Relationships between PNC at necropsy and comorbidities were examined by comparing medians (Q1, Q3) for NHPs with and without each comorbidity, and the Mann-Whitney U test was used to compare the distributions of PNC at necropsy based on comorbidity presence. In **Table 5**, cardiac fibrosis was the only measured comorbidity with a significant difference in PNC; the high cardiac fibrosis group had a median PNC of 0.65 ng/mL at necropsy, compared to 0.38 ng/mL for the low cardiac fibrosis group, and these PNC distributions were significantly different (p = 0.007). In sum, these results demonstrate that, despite multi-organ injury caused by TBI, serum PNC is a marker of cardiac fibrosis of irradiated NHPs.

**Table 5.** Relationships between PNC and comorbidities at necropsy.

| Comorbidity | Absent |  | Present |  | p-value |
| --- | --- | --- | --- | --- | --- |
|  | N | Median PNC (ng/mL)<br>(Q1, Q3) | N | Median PNC (ng/mL)<br>(Q1, Q3) |  |
| Hypertension | 39 | 0.40 (0.29, 0.62) | 17 | 0.45 (0.23, 0.77) | 0.803 |
| Overweight | 37 | 0.37 (0.28, 0.48) | 19 | 0.62 (0.39, 0.72) | 0.072 |
| Underweight | 43 | 0.40 (0.29, 0.67) | 13 | 0.43 (0.30, 0.48) | 0.801 |
| Tumor (carcinoma, sarcoma, both) | 38 | 0.35 (0.24, 0.66) | 18 | 0.49 (0.39, 0.74) | 0.120 |
| Gastrointestinal | 38 | 0.44 (0.29, 0.66) | 18 | 0.39 (0.26, 0.64) | 0.488 |
| Heart murmur | 31 | 0.39 (0.29, 0.65) | 25 | 0.41 (0.23, 0.68) | 0.961 |
| Lung | 38 | 0.40 (0.26, 0.66) | 18 | 0.44 (0.31, 0.64) | 1.000 |
| Diabetes | 40 | 0.40 (0.29, 0.67) | 16 | 0.57 (0.22, 0.64) | 0.790 |
| Cataracts | 21 | 0.36 (0.30, 0.68) | 35 | 0.41 (0.24, 0.63) | 0.710 |
| Hepatic cysts | 44 | 0.39 (0.28, 0.63) | 12 | 0.58 (0.39, 0.72) | 0.122 |
| Kidney | 32 | 0.39 (0.24, 0.55) | 24 | 0.47 (0.32, 0.71) | 0.145 |
| Arthritis | 15 | 0.37 (0.27, 0.49) | 41 | 0.43 (0.30, 0.67) | 0.350 |
| Dermatitis | 32 | 0.39 (0.25, 0.52) | 24 | 0.51 (0.32, 0.69) | 0.289 |
| Testicular atrophy | 21 | 0.38 (0.24, 0.48) | 35 | 0.45 (0.29, 0.69) | 0.163 |
| Irradiated | 10 | 0.37 (0.23, 0.47) | 46 | 0.42 (0.29, 0.67) | 0.252 |
| High cardiac fibrosis (2-3) | 44 | 0.38 (0.25, 0.52) | 12 | 0.65 (0.52, 0.79) | 0.007** |
| High pulmonary fibrosis (2-3) | 45 | 0.39 (0.28, 0.67) | 11 | 0.49 (0.43, 0.66) | 0.343 |

P-values are obtained from Mann-Whitney U tests comparing the distributions of PNC values at necropsy when the specified comorbidity is absent or present (ever). Counts (N) represent the count of NHPs with non-missing PNC information at necropsy. Lung comorbidity was defined by abnormal CT density. Kidney comorbidity was defined by cyst, nephromegaly, abnormal CT density, or abnormal blood urea nitrogen (BUN) levels. Criteria for diagnosis of comorbidities has previously been reported<sup>42</sup>. (\* $p < 0.05$ ; \*\* $p < 0.01$ ).

## DISCUSSION

Exposure to TBI is deleterious to all organ systems and often leads to comorbidities associated with or characterized by tissue fibrosis. Indeed, NHPs from this long-term survivor cohort carry comorbidities such as diabetes, hypertension, tumors, gastrointestinal disease, kidney disease, liver disease, and heart disease^27,28^. It is noteworthy that despite multi-organ injury from TBI, PNC remains resilient to confounding comorbidities as a marker of cardiac fibrosis and diastolic echocardiogram parameters. However, the ideal timing for using serum PNC for RRHD detection or prediction remains to be established. Furthermore, while this study suggests organ-specificity for diagnostic purposes in RRHD, our previous studies allude to PNC as a biomarker of heart failure from multiple etiologies^22–24^.

In the present study, irradiated NHPs with severe cardiac fibrosis exhibited significantly higher ejection fractions but also significantly elevated diastolic filling pressures compared to non-irradiated animals with similar fibrosis scores. These findings support existing evidence that radiation exposure to the heart more commonly results in diastolic dysfunction, even when ejection fraction remains preserved. One possible explanation is that, although the severity of fibrosis appears comparable between irradiated and non-irradiated animals with high cardiac fibrosis scores, differences in the distribution of fibrosis, collagen content, and collagen cross-linking may underlie the observed functional divergence^46–49^. On the contrary, echocardiogram data from unirradiated NHPs with high cardiac fibrosis scores were limited, and this observation could be due to a limited sample set.

Echocardiography is a non-invasive surrogate for measuring cardiac remodeling due to cardiac injury, stress, or disease. Low or high early E to late A ventricular filling velocities can indicate stiffness in the ventricular myocardium due to fibrotic remodeling. Elevated E/e’ values indicate increased left ventricular filling pressures, which are also often due to fibrotic remodeling of the ventricle. Increased stiffness and prolonged exposure to elevated ventricle filling pressures during diastole lead to enlargement of the left atrium, increased LA volume, and LAA. Many of the echocardiogram parameters used in assessing diastolic dysfunction significantly correlate with serum PNC levels in these NHPs. Serum PNC levels could potentially be used as a surrogate measure of diastolic dysfunction. Several papers have examined the association between serological BNP/NT-proBNP and cardiotoxicity related to RT^21,50–54^. For example, in a study using a cohort of long-term breast cancer survivors who received RT only, the changes in BNP levels were significantly higher in patients with increased cardiac and left ventricular radiation doses^51^. In another study including patients treated with thoracic RT, the percentage of patients with ECG changes among patients with elevated NT-proBNP levels was significantly higher than that among patients without elevated NT-proBNP levels during the 6 to 12 months follow-up time. Studies evaluating traditional cardiac biomarkers in RRHD in Rhesus macaques are limited. One study demonstrated the utility of cardiac troponin I (cTnI) and NT-proBNP in identifying hypertrophic cardiomyopathy and other cardiac diseases in NHPs; however, these animals were not irradiated. Both cTnI and NT-proBNP had significant correlations with various echocardiogram parameters in this study^55^. Although BNP/NT-proBNP serves the primary role of ruling out HF in emergency point-of-care settings and guiding treatment within the HF patient population^19^, the unavoidable confounding variables and comorbidities found within the general population contribute to the lack of sensitivity or specificity of BNP/NT-proBNP as a biomarker for assessing the risk of HF^56–58^. However, our prior work demonstrated that serological PNC levels are not affected by major confounders such as age, sex, and BMI,^22^ suggesting that PNC could offer advantages over BNP/NT-proBNP in predicting the risk of heart disease following radiation exposure.

Together, our findings demonstrate the significant utility of serum PNC for identifying cardiac fibrosis and significant correlations to echocardiogram parameters of diastolic dysfunction in RRHD in Rhesus macaques. These findings suggest the promise of developing serum PNC as a biomarker of RRHD among cancer patients who receive thoracic radiotherapy and survivors of acute radiation exposure from nuclear accidents.

## LIMITATIONS

The inferences derived from this retrospective study are limited by the radiation exposure characteristics. Positive aspects of the exposure strategy include uniformity of the exposure field and well-controlled dose and dose rate. However, the uniformity of the exposure, while providing experimental rigor and reproducibility, would be unlikely in the case of a real-world exposure. The total radiation dose was lower than that for thoracic radiotherapy, and the dose was not fractionated. Relative to nuclear explosive devices, the total dose was higher than that experienced by survivors, and consisted of photons only, not the mixed photon-proton-neutron spectrum likely in the case of a nuclear device. In addition, our results cannot exclude the possibility that fibrosis in organs other than the heart and lungs could potentially contribute to elevated serum PNC because not every organ was scored for fibrosis. While echocardiographic parameters provide a reasonable non-invasive assessment, invasive hemodynamic measurements remain the gold standard for evaluating diastolic filling pressures. Moreover, echocardiography was not performed in parallel with serum sample collection; therefore, correlations were conducted on echocardiogram measurements within 6 months of serum sample collection. The cohort consisted of only male Rhesus macaques. Serum samples from the unirradiated sub-cohorts were limited. Lastly, conventional echocardiogram threshold values defining diastolic dysfunction for Rhesus macaques have not been adequately defined in the literature^59^.

## PERSPECTIVES

### Competency in Medical Knowledge

Radiation-related heart disease can be an undetected side effect of radiation therapy, especially in individuals undergoing treatment for cancers situated close to the chest, and has drawn interest in recent years. Yet, knowledge of its underlying mechanisms, detection, and effective treatment options remains incomplete. Identifying novel biomarkers for evaluating risk is vital for mitigating the effects of radiation exposure.

### Translational Outlook

This study sets the stage for a continued effort to define qualifying biomarkers for FDA approval for evaluating radiation-related heart disease (RRHD). Future studies involving planned clinical trials with the inclusion of pro-N-cadherin (PNC) and other biomarkers that utilize more clinically relevant radiation exposure protocols are critical for developing PNC as a biomarker of RRHD.

## Acknowledgment

We thank Dr. Eva Guinan for her constructive comments. The funding sources for this manuscript are: National Institutes of Health U01AI189426 (SVP and C-LL), U01AI50578 (JMC), U19AI67798 (JMC), 5P30-CA014236-50 (DN), DOD/CDMRP award W81XWH-15-1-0574 (JMC and TCR), and Duke University School of Medicine Whitehead Scholar Award (C-LL).

## Author Contributions Statement

PDF, SVP, and C-LL conceptualized the PNC project; PDF, GOD, GWS, JDO, KMO, KTM, TCR, DK, C-LL designed and performed experiments; TCR led the echo measures focusing on diastolic function; PDF and DN conducted statistical analysis; TCR, JMC, SVP, and C-LL acquired funding; TCR, JMC, SVP, and C-LL supervised the project; PDF, DN and C-LL wrote the original draft; All authors reviewed and edited the final manuscript; SVP, and C-LL contributed equally to this manuscript.

## ABBREVIATIONS AND ACRONYMS

RRHD: radiation-related heart disease
PNC: pro-N-cadherin
HF: heart failure
NHP: non-human primate
TBI: total-body irradiation
RT: radiation therapy
BNP: B-type natriuretic peptide
RLEC: Radiation Late Effects Cohort

