## Supplementary Tables for "Serum pro-N-cadherin: a biomarker of cardiac fibrosis and diastolic dysfunction in irradiated non-human primates"

**Supplemental Appendix**

Serum pro-N-cadherin: a biomarker of cardiac fibrosis and diastolic dysfunction in the radiation late effects cohort of non-human primates

Supplemental Table 1. Echocardiogram summary between sub-cohorts, 30 months from necropsy……………..........................................................................................................Page 2

Supplemental Table 2. Echocardiogram summary between sub-cohorts, 12 months from necropsy……………..........................................................................................................Page 4

Supplementary Table 3. Echocardiogram relationships with cardiac fibrosis and PNC, after adjustment for BSA and age……………...........................................................................Page 6

Supplementary Figure 1. NHP plasma serial dilution fits non-linear regression ……………...................................................................................................................…..Page 7

Supplementary Figure 2. Distribution of echocardiograms performed within 6 months of serum collection by sub-cohort…………………...........................................................................Page 8

**Supplemental Table 1. Echocardiogram summary between sub-cohorts, 30 months from necropsy**

| | NO IR F0-1 | NO IR F2-3 | IR F0-1 | IR F2-3 |
|---|---|---|---|---|
| **EF A4C** | 69.19 (54.97, 74.26) * N=12 | 52.08 (49.31, 60.58) * €€€ ¥¥ N=10 | 67.31 (63.16, 71.70) €€€ N=72 | 70.06 (60.64, 72.87) ¥¥ N=17 |
| **EF BP** | 71.87 (64.08, 74.06) N=11 | 60.89 (54.24, 64.70) € ¥ N=6 | 69.92 (66.77, 73.64) € N=52 | 72.98 (62.43, 76.97) ¥ N=9 |
| **DIFD** | 10.74 (6.70, 15.09) Ω ‡ N=12 | 7.14 (4.60, 10.78) €€€ ¥¥¥ N=10 | 21.48 (14.50, 28.50) Ω €€€ N=71 | 24.45 (20.02, 28.17) ‡ ¥¥¥ N=17 |
| **A' LATERAL** | 8.11 (6.87, 9.04) N=12 | 5.64 (4.27, 7.59) N=10 | 7.19 (5.69, 9.02) N=73 | 6.81 (5.75, 8.85) N=18 |
| **A' SEPTAL** | 7.29 (6.20, 7.80) N=7 | 5.77 (5.11, 6.77) N=7 | 5.87 (4.90, 8.44) N=47 | 6.14 (6.07, 7.03) N=11 |
| **E' LATERAL** | 12.04 (9.37, 12.88) N=12 | 11.62 (6.22, 13.47) N=10 | 10.17 (9.04, 11.99) N=73 | 9.47 (7.93, 10.34) N=18 |
| **E/E' LATERAL** | 6.42 (5.77, 8.87) ΩΩ ‡‡‡ N=12 | 7.87 (6.26, 10.35) ¥¥ N=10 | 9.99 (8.68, 11.29) ΩΩ §§ N=73 | 12.25 (11.39, 14.83) ‡‡‡ ¥¥ §§ N=17 |
| **E'/A' LATERAL** | 1.36 (1.23, 1.54) N=12 | 1.41 (1.18, 1.95) N=10 | 1.45 (1.15, 1.79) N=73 | 1.24 (1.01, 1.75) N=18 |
| **E/A** | 1.10 (0.97, 1.23) N=12 | 1.12 (0.98, 1.29) N=10 | 1.24 (1.09, 1.47) N=93 | 1.15 (0.93, 1.74) N=17 |
| **MV E VEL** | 0.79 (0.63, 0.82) ΩΩ ‡‡ N=12 | 0.84 (0.74, 0.92) €€€ N=10 | 1.01 (0.93, 1.17) ΩΩ €€€ N=93 | 1.09 (0.91, 1.29) ‡‡ N=17 |

| | | | | |
|---|---|---|---|---|
| **LA VOL A2C** | 3.37 (2.55, 4.65)<br><br>N=12 | 4.99 (3.17, 6.12)<br>€€€<br>N=10 | 2.26 (1.71, 2.97)<br>€€€<br>N=73 | 2.69 (1.73, 3.30)<br><br>N=17 |
| **LA VOL BP** | 4.05 (3.19, 5.13)<br><br>N=12 | 5.67 (3.93, 6.16)<br>€€€<br>N=10 | 3.11 (2.41, 3.69)<br>€€€<br>N=72 | 3.68 (3.13, 4.56)<br><br>N=16 |
| **LAA A2C** | 2.89 (2.51, 3.49)<br>Ω<br>N=12 | 3.54 (2.88, 3.92)<br>€€€ ¥<br>N=10 | 2.12 (1.70, 2.60)<br>Ω €€€<br>N=73 | 2.50 (1.71, 2.70)<br>¥<br>N=17 |
| **LA DIAMETER SYSTOLE** | 17.76 (16.62, 19.65)<br>ΩΩ<br>N=12 | 19.72 (17.40, 22.42)<br>€€€<br>N=10 | 14.36 (13.00, 17.12)<br>ΩΩ €€€ §§<br>N=97 | 17.14 (15.44, 19.35)<br>§§<br>N=22 |
| **LA/AO SYSTOLE** | 1.62 (1.36, 1.82)<br><br>N=12 | 1.67 (1.48, 1.73)<br><br>N=10 | 1.54 (1.35, 1.71)<br>§§<br>N=71 | 1.80 (1.62, 2.00)<br>§§<br>N=17 |

EF A4C = Left ventricular ejection fraction apical 2-chamber, EF BP = Left ventricular ejection fraction biplane, DIFD = Left ventricle - major axis length diastolic difference - end diastole, a' lateral = Left ventricle peak diastolic tissue velocity during atrial systole – lateral mitral annulus, a' septal = Left ventricle peak diastolic tissue velocity during atrial systole – septal mitral annulus, e' lateral = Left ventricular peak early diastolic tissue velocity – lateral mitral annulus, E/e' lateral = ratio of mitral valve peak velocity to left ventricle peak tissue velocity E-Wave, e'/a' lateral = ratio of the peak early to late lateral mitral annular filling velocity, E/A = ratio of the peak early to late transmitral flow velocity, MV E vel = Mitral valve E wave peak velocity, LA vol A2C = Left atrial volume apical 2-chamber – end systole, LA vol BP = Left atrial volume biplane – end systole, LAA A2C = Left atrial area apical 2-chamber – end systole, LA diameter systole = Left atrium diameter systole – antero-posterior, LA/AO systole = Left atrium diameter to aortic root diameter – systole. Counts (N) represent total echocardiogram examinations for each measure with corresponding serum samples within 6 months of examination over 0-2 years prior to necropsy. Median (Q1, Q3) values are presented. P-values were calculated from one-way Kruskal-Wallis tests across strata for each measure, with Dunn's post-hoc tests (* p < 0.05; ** p < 0.01; *** p < 0.001), (€ p < 0.05; €€ p < 0.01; €€€ p < 0.001), (¥ p < 0.05; ¥¥ p < 0.01; ¥¥¥ p < 0.001), (Ω p < 0.05; Ω Ω p < 0.01; Ω Ω Ω p < 0.001), (‡ p < 0.05; ‡‡ p < 0.01; ‡‡‡ p < 0.001), (§ p < 0.05; §§ p < 0.01; §§§ p < 0.001).

**Supplemental Table 2. Echocardiogram summary between sub-cohorts, 12 months from necropsy**

| | IR F0-1 | IR F2-3 |
|---|---|---|
| **EF A4C** | 68.21 (64.06, 74.52)<br>N=22 | 69.47 (58.60, 72.66)<br>N=7 |
| **EF BP** | 70.05 (67.08, 75.06)<br>N=21 | 72.26 (60.08, 73.58)<br>N=7 |
| **DIFD** | 22.65 (16.15, 30.61)<br>N=22 | 25.70 (20.57, 30.08)<br>N=7 |
| **A' LATERAL** | 7.72 (6.57, 9.47)<br>N=22 | 8.42 (5.96, 10.22)<br>N=8 |
| **A' SEPTAL** | 7.92 (5.13, 9.01)<br>N=20 | 6.67 (6.41, 7.03)<br>N=3 |
| **E' LATERAL** | 10.71 (9.51, 11.78)<br>N=22 | 9.70 (8.46, 10.26)<br>N=8 |
| **E/E' LATERAL** | 9.96 (8.90, 11.12)<br>***<br>N=22 | 12.47 (11.85, 14.56)<br>***<br>N=8 |
| **E'/A' LATERAL** | 1.37 (1.18, 1.63)<br>N=22 | 1.09 (0.76, 1.48)<br>N=8 |
| **E/A** | 1.13 (0.98, 1.25)<br>N=31 | 1.00 (0.90, 1.95)<br>N=8 |
| **MV E VEL** | 0.99 (0.91, 1.16)<br>N=31 | 1.19 (0.99, 1.40)<br>N=8 |
| **LA VOL A2C** | 2.30 (2.19, 2.89)<br>N=22 | 2.49 (1.91, 3.61)<br>N=7 |
| **LA VOL BP** | 3.00 (2.69, 3.47)<br>N=21 | 3.68 (2.66, 4.34)<br>N=6 |

| | | |
|---|---|---|
| **LAA A2C** | 2.20 (2.06, 2.57)<br><br>N=22 | 2.30 (1.84, 2.92)<br><br>N=7 |
| **LA DIAMETER SYSTOLE** | 14.42 (13.40, 17.00)<br>**<br><br>N=32 | 18.40 (15.85, 20.50)<br>**<br><br>N=9 |
| **LA/AO SYSTOLE** | 1.35 (1.22, 1.63)<br>*<br><br>N=21 | 1.80 (1.68, 1.97)<br>*<br><br>N=7 |

EF A4C = Left ventricular ejection fraction apical 2-chamber, EF BP = Left ventricular ejection fraction biplane, DIFD = Left ventricle - major axis length diastolic difference - end diastole, a' lateral = Left ventricle peak diastolic tissue velocity during atrial systole – lateral mitral annulus, a' septal = Left ventricle peak diastolic tissue velocity during atrial systole – septal mitral annulus, e' lateral = Left ventricular peak early diastolic tissue velocity – lateral mitral annulus, E/e' lateral = ratio of mitral valve peak velocity to left ventricle peak tissue velocity E-Wave, e'/a' lateral = ratio of the peak early to late lateral mitral annular filling velocity, E/A = ratio of the peak early to late transmitral flow velocity, MV E vel = Mitral valve E wave peak velocity, LA vol A2C = Left atrial volume apical 2-chamber – end systole, LA vol BP = Left atrial volume biplane – end systole, LAA A2C = Left atrial area apical 2-chamber – end systole, LA diameter systole = Left atrium diameter systole – antero-posterior, LA/AO systole = Left atrium diameter to aortic root diameter – systole. Counts (N) represent total echocardiogram examinations with corresponding serum samples for each measure within 6 months of examination over 12 months prior to necropsy. Median (Q1, Q3) values are presented. P-values were calculated from Mann-Whitney U tests across strata for each measure. (* $p < 0.05$; ** $p < 0.01$; *** $p < 0.001$).

**Supplementary Table 3. Echocardiogram relationships with cardiac fibrosis and PNC, after adjustment for BSA and age**

In Table 2 we observed key echocardiogram measures which display significant relationships with irradiation status and cardiac fibrosis. In Table 4 we observed key echocardiogram measures which display significant correlations with PNC. To adjust these measures for age at echocardiogram measurement and body surface area (BSA), we explored these relationships within multivariable regression frameworks. This analysis was restricted to the closest measurement to necropsy for each NHP, limited to 12 months prior, and due to small sample sizes, only within the irradiated group. The first framework explored the use of each echocardiogram measure as the dependent variable, with high cardiac fibrosis as the independent variable, additionally adjusted for BSA and age. The second framework explored the use of each echocardiogram measure as the dependent variable, with PNC as the independent variable, additionally adjusted for BSA and age. Some echocardiogram measures display significant relationships with high cardiac fibrosis, and for others, statistical significance was observed with PNC. In each regression framework, E/e' lateral had significant positive associations with high cardiac fibrosis and PNC, and this significance remained after adjustments for BSA and age.

| Echocardiogram linear regressions with cardiac fibrosis and PNC, with adjustment for BSA and age | | | | |
|---|---|---|---|---|
| | Linear regression; using high cardiac fibrosis (F2-3 vs. F0-1) to predict key echocardiogram measure | | Linear regression; using PNC to predict key echocardiogram measure | |
| | β (95% CI) coefficient of high cardiac fibrosis | | β (95% CI) coefficient of PNC | |
| Echocardiogram measure | Unadjusted | Adjusted for BSA and age | Unadjusted | Adjusted for BSA and age |
| E/e' lateral | **3.50 (1.68, 5.33)** | **3.29 (1.11, 5.48)** | **4.06 (1.90, 6.22)** | **3.99 (1.22, 6.75)** |
| LA diameter systole | **3.40 (1.25, 5.55)** | **4.16 (1.60, 6.72)** | 1.67 (-1.20, 4.54) | 1.48 (-1.90, 4.86) |
| LA/AO systole | **0.36 (0.08, 0.63)** | **0.43 (0.11, 0.75)** | 0.106 (-0.25, 0.46) | 0.10 (-0.35, 0.54) |
| AV PGMax | -0.24 (-2.21, 1.72) | 0.89 (-1.26, 3.04) | **-2.60 (-4.65, -0.55)** | -2.06 (-4.66, 0.53) |
| Confidence intervals (CI)s from Wald technique, highlighted in bold when statistical significance is achieved ($p < 0.05$). | | | | |

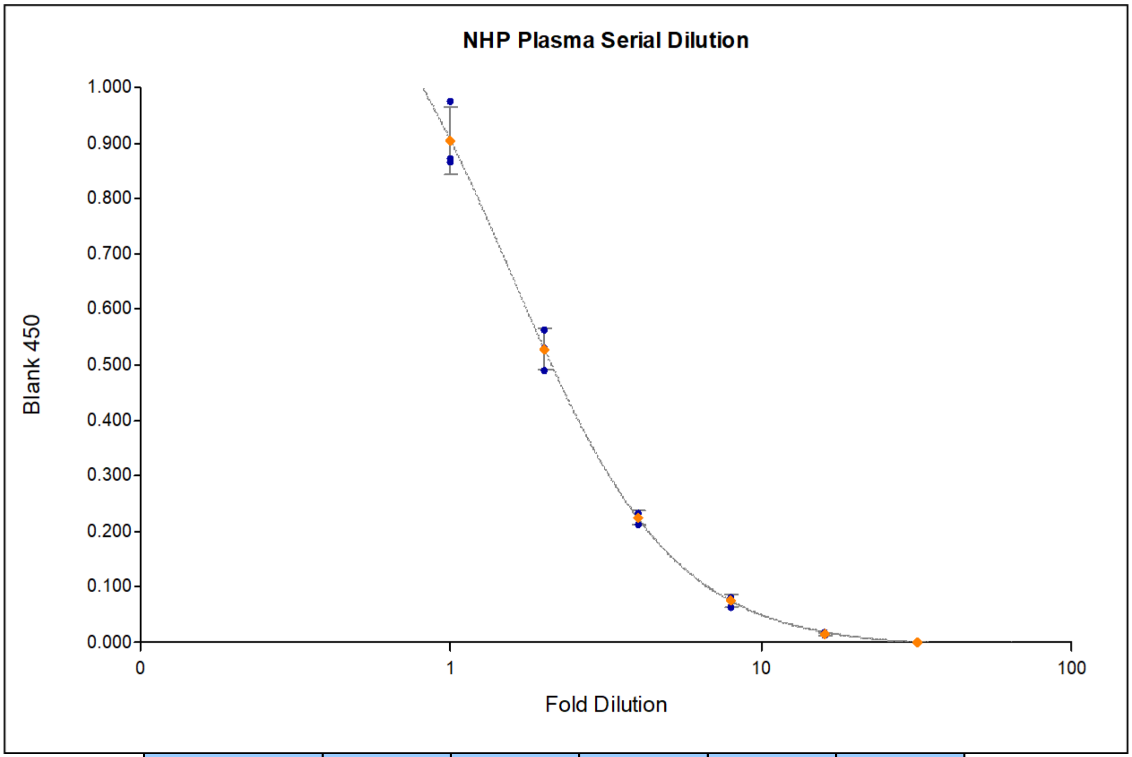


| Curve Formula | A | B | C | D | R2 |
|---|---|---|---|---|---|
| Y = (A-D)/(1+(X/C)^B) + D | 1.34 | 1.67 | 1.57 | -0.00894 | 1 |

**Supplementary Figure 1. NHP plasma serial dilution fits non-linear regression.** The pooled NHP plasma sample was serially diluted in triplicate and quantified by PNC ELISA using previously established methods ($r^2 = 1.00$). Dilutions started with neat (1-fold), followed by 2-fold, 4-fold, 8-fold, 16-fold, and 32-fold.

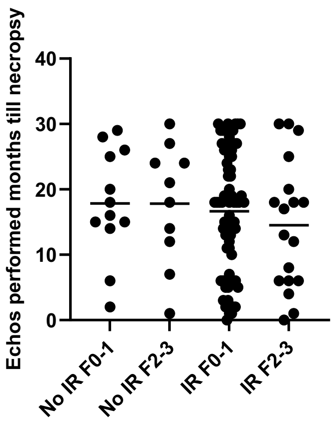


**Supplementary Figure 2. Distribution of echocardiograms performed within 6 months of serum collection by sub-cohort.** The distribution of time of echocardiograms prior to necropsy was not significantly different between any of the sub-cohorts. (No IR F0-1 vs. No IR F2-3, p>0.999; No IR F0-1 vs. IR F0-1, p=0.976; No IR F0-1 vs. IR F2-3, p=0.768; No IR F2-3 vs. IR F0-1, p=0.983; No IR F2-3 vs. IR F2-3, p=0.802; IR F0-1 vs. IR F2-3, p=0.815)
